# Predicting the unseen: a diffusion-based debiasing framework for transcriptional response prediction at single-cell resolution

**DOI:** 10.1101/2025.09.12.675662

**Authors:** Ergan Shang, Yuting Wei, Kathryn Roeder

## Abstract

Predicting cellular responses to genetic perturbations is critical for advancing our understanding of gene regulation. While single-cell CRISPR perturbation assays such as Perturb-seq provide direct measurements of gene function, the scale of these experiments is limited by cost and feasibility. This motivates the development of computational approaches that can accurately infer responses to unmeasured perturbations from related experimental data. We introduce dbDiffusion, a generative framework that integrates diffusion models with classifier-free guidance derived from perturbation information, operating in latent space through a variational autoencoder (VAE). Diffusion models are probabilistic generative models that approximate data distributions by reversing a Markovian diffusion process, progressively denoising Gaussian noise into structured outputs. By exploiting biological similarities in gene expression profiles and relationships among perturbations, dbDiffusion enables the conditional generation of gene expressions for previously unobserved perturbations. In contrast to competing approaches, dbDiffusion does not rely on LLM or foundation models, which have been found to yield unsatisfactory results. Rather, it leverages embeddings derived from measured perturbations to generalize to unseen perturbations, effectively transferring information across related experimental conditions. In benchmarking against state-of-the-art methods on Perturb-seq datasets, dbDiffusion demonstrates superior accuracy in predicting perturbation responses. A methodological innovation of dbDiffusion is the integration of prediction-powered inference, which corrects for biases inherent in generative models and enables statistically rigorous downstream tasks, including identification of differentially expressed genes. By combining deep generative modeling with principled inference, dbDiffusion establishes a scalable computational framework for predicting and analyzing transcriptomic perturbation responses, significantly extending the utility of Perturb-seq experiments.

## 1 Introduction

Despite extensive efforts in functional genomics, the roles of many genes remain poorly characterized. While sequence-based annotations offer broad predictions of molecular function, the specific mechanisms, biological pathways, and cellular contexts in which genes operate are often unclear (Juncker et al., 2009). The integration of pooled genetic perturbations with single-cell RNA sequencing -exemplified by Perturb-seq (Dixit et al., 2016; Adamson et al., 2016) and CROP-seq (Jaitin et al., 2016; Datlinger et al., 2017) now enables systematic high-resolution dissection of gene function. By combining CRISPR-based perturbations with transcriptional profiling in single cells, these approaches reveal how targeted genetic changes shape cellular states and regulatory programs. Such tools transform our ability to map gene function, decode regulatory networks (Rubin et al., 2019; Geiger-Schuller et al., 2023), and link genetic variation to phenotype with unprecedented precision Barrangou and Doudna (2016); Frangieh et al. (2021); Schnitzler et al. (2024).

High-throughput technologies have greatly expanded our ability to profile the effects of genetic perturbations, but the scale of possible targets precludes comprehensive experimental screening (Yao et al., 2024). Computational methods are critical for prioritizing perturbations, yet existing models often struggle to provide satisfactory results. There is a pressing need for predictive frameworks that generalize to unseen perturbations and guide experimental design with greater accuracy and efficiency (Peidli et al., 2024; Yang et al., 2025).

Recent methods—including GEARS (Roohani et al., 2024), scLAMBDA (Wang et al., 2024), and singlecell foundation models such as scGPT (Cui et al., 2024)—have aimed to predict perturbation effects for previously unobserved genes. However, despite their promise, deep learning-based models have yet to consistently outperform simple baseline approaches (Ahlmann-Eltze et al., 2025; Csendes et al., 2025; Wong et al., 2025), highlighting the need for more robust and generalizable predictive frameworks.

To overcome limitations of current predictive models, we introduce *debiased diffusion (dbDiffusion)*, a deep generative framework that leverages a latent diffusion model with classifier-free guidance, combined with a customized embedding strategy, to predict single-cell responses to genetic perturbations. ^1^ A key feature of the method is the debiasing step, which takes advantage of the embedding in a 2-step process to improve the predicted quantities. While other methods, such as scLAMBDA, aim to generate realistic outcomes for a sample of cells, dbDiffusion aims to produce an accurate estimate of the mean effect on each gene of an unseen perturbation, along with a quantification of uncertainty through a confidence interval on the effect size. Across multiple benchmark datasets, dbDiffusion consistently outperforms existing approaches.

## 2 Results

### 2.1 Methods Overview

The dbDiffusion framework, adapted from cfDiffusion Zhang et al. (2025), consists of two main components: an autoencoder (AE) and a diffusion model (Figure 1). The AE compresses the high-dimensional input data into a latent representation that preserves the biological signal. These latent features serve as input to the diffusion model. To implement classifier-free guidance, we estimate an embedding for each perturbation, which steers latent feature generation toward the distribution of the target class. After training, the diffusion model can synthesize latent representations from random noise conditioned on a specified perturbation type. The AE decoder subsequently reconstructs the corresponding gene expression profiles from these generated latent features.

**Figure 1.**
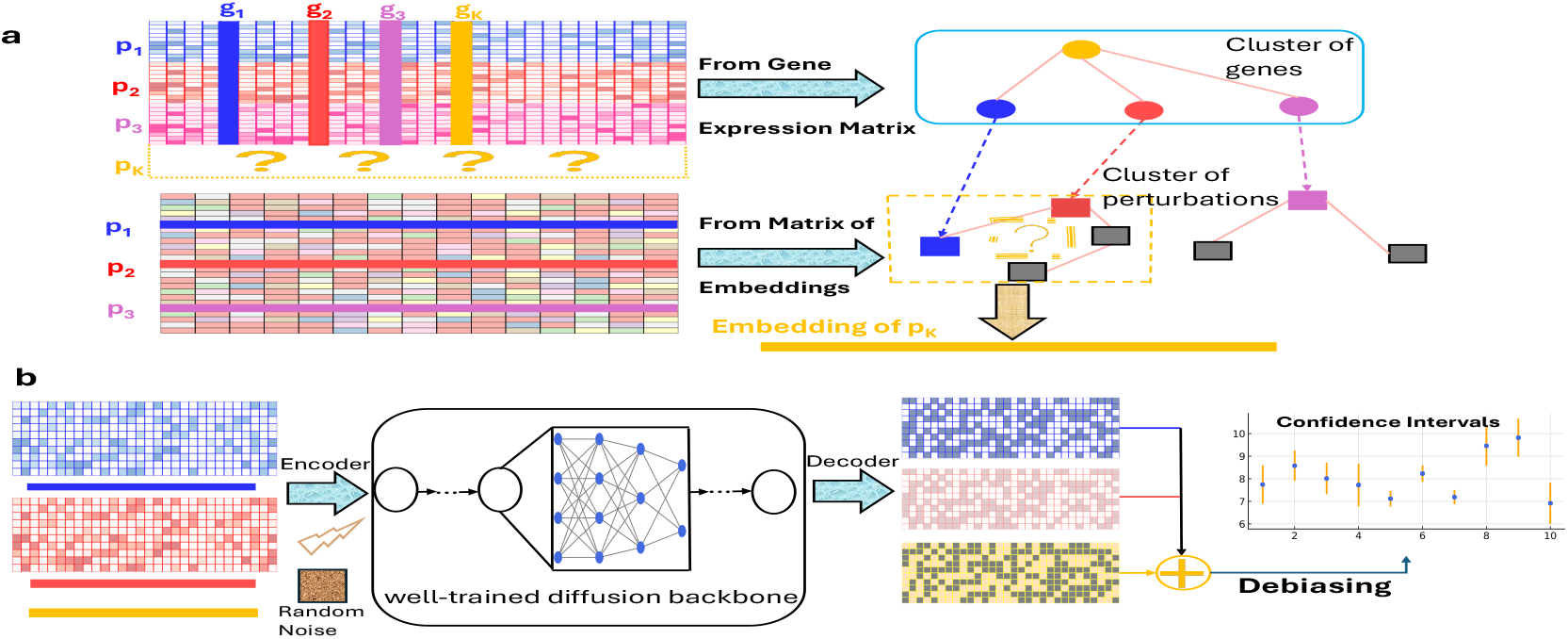
Overview of dbDiffusion. dbDiffusion takes single-cell genetic perturbation datasets along with the embeddings of each perturbation, which can be calculated by PCA of gene expression matrix or effect size matrix, for example, as input. dbDiffusion will first generate the embedding of the perturbation to be evaluated using both clusters of genes and clusters of perturbations observed. After mapping those genes in the same cluster of the perturbation targeted into the cluster of perturbations, the embedding is calculated after the best cluster of perturbation is selected. With embeddings of all perturbations in the best cluster, dbDiffusion uses the diffusion model that has been well-trained on the genetic perturbation dataset to generate as many samples as possible. After debiasing, dbDiffusion makes inference on the mean of gene expression by constructing confidence intervals.

### 2.2 Diffusion model for Tissue Data

Using a compendium of single-cell transcriptomic data from the musculus model organism, comprising 12 distinct tissues (Tabula Muris Consortium et al., 2018), we illustrate how an embedding can be empirically derived and cells can be generated using our classifier-free diffusion model. Based on the original musculus dataset containing 18,996 genes, we first select 2,000 highly variable genes and perform pseudo-bulk aggregation by tissue type. We then extract the top three principal components, which serve as a low-dimensional tissue embedding for visualization in Figure 2a and for demonstration in Figure 2d. In line with cfDiffusion Zhang et al. (2025), which trains on the full, original dataset, we also train our classifier-free diffusion model on the full dataset, but incorporate a higher-dimensional (64D) continuous PC-based embedding (Figure 2c) instead of one-hot labels (Figure 2b). Note that the cfDiffusion method does not predict as well, as it only permits a fixed set of discrete one-hot embeddings corresponding to observed categories. In contrast, our model accepts a continuous embedding representation, enabling interpolation and meaningful predictions beyond the training categories. Finally, using only knowledge that the spleen lies close to the lung and marrow tissues, we generate spleen cells using an interpolation of the lung and marrow embedding. Again, the generated spleen cells match the targeted mean expression well, but fail to capture the full variability of the original cells (Figure 2d). Provided that our goal is to determine the mean gene expression of a particular class, the performance of this generator is sufficient.

**Figure 2.**
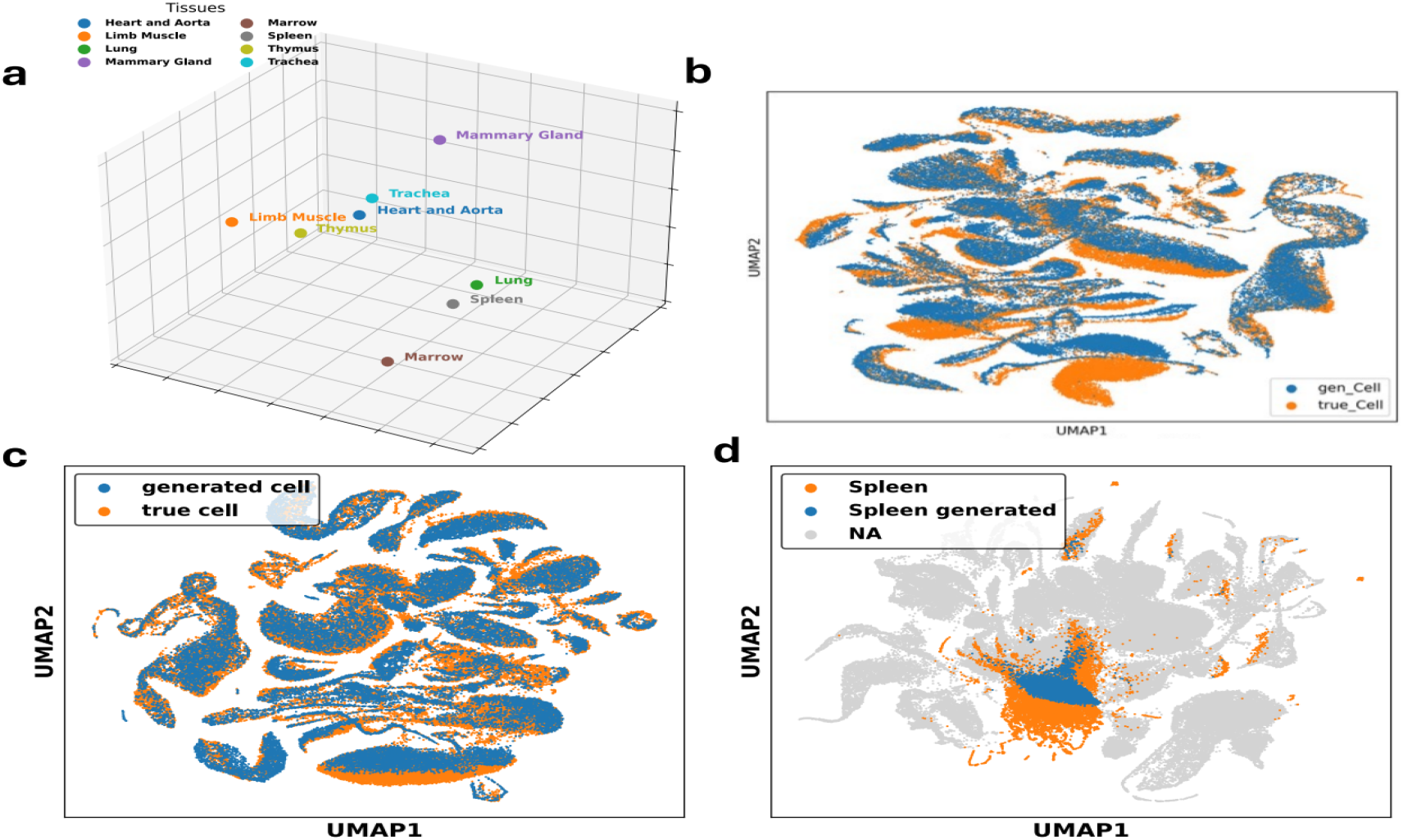
**a**. Illustration of the three-dimensional embeddings of different cell types. Only tissue types of interest are shown. **b**. UMAP visualization of generated and true cells across all cell types, taken from cfDiffusion (Zhang et al., 2025). **c**. UMAP visualization between generated cells and the true cells across all cell types using the classifier-free diffusion model with embeddings of 64 dimensions obtained by PCA. **d**. UMAP visualization highlighting the true Spleen cells and generated Spleen cells, which are obtained using embedding as the linear combination of the three-dimensional embedding of Marrow and Lung in **a**. in the classifier-free diffusion model.

### 2.3 Preliminaries for diffusion models

A diffusion model typically encompasses two Markov processes: a forward process and a reverse process. In the forward process, one progressively injects noise into the data samples to diffuse the data, and produces a sequence of *d*-dimensional random vectors *X*_1_ → *X*_2_ → …→ *X*_*T*_ as follows:

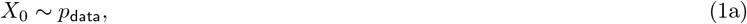

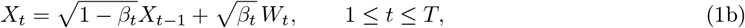

where {*W*_1≤*t*≤*T*_} denotes a sequence of independent noise vectors drawn from 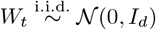. Taking the continuum limit of (1), the process satisfies the stochastic differential equation (SDE):

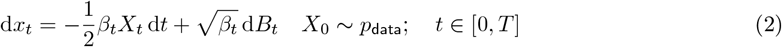

for some function *β*_*t*_ : [0, *T*] → ℝ, where (*B*_*t*_)_*t*∈[0,*T*]_ is a standard Brownian motion in ℝ^*d*^. However, in the reverse process, the reverse chain *Y*_*T*_ → *Y*_*T* −1_ → … → *Y*_1_ is designed to (approximately) revert the forward process, allowing one to transform pure noise *Y*_*T*_ ∼ 𝒩 (0, *I*_*d*_) into new samples with matching distributions as the original data. The classical SDE theory (Anderson, 1982; Haussmann and Pardoux, 1986) guarantees that its time reversal 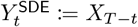 satisfies:

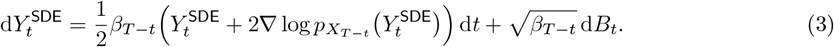

Here, 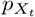 denotes the marginal distribution of *X*_*t*_ in the forward SDE (2) and 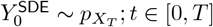.

From the continuous perspective, it is clear that if the score function 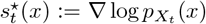 were known exactly, the reverse process could be uniquely identified and thus sampling would be easy. However, in practice, the score functions need to be learned from training samples. Suppose that we are given score estimates 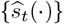 at *t* = 1, …, *T* . Among other algorithms, the renowned (Denoising Diffusion Probabilistic Model) DDPM algorithm Ho et al. (2020) serves as a stochastic sampler that recursively generates samples via

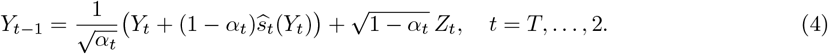

with *Y*_T_ ∼ 𝒩 (0, *I*_d_). Here, 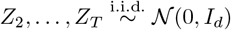 is a sequence of i.i.d. standard Gaussian random vectors in ℝ^*d*^ that is independent of 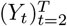.

For many practical tasks, a natural goal is to generate samples given some constraints or contexts, which motivates the study of *conditional* diffusion models. In a nutshell, conditional diffusion models aim to generate samples from the distribution *p*(*x* | *c*) conditional on a specific label or context. In order to do so, it requires estimating the conditional score, which according to Bayes’ rule satisfies

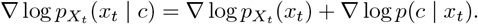

A prominent scheme that enjoys tremendous practical success is the so-called classifier-free guidance Ho and Salimans (2022), which approximates the conditional score by the following

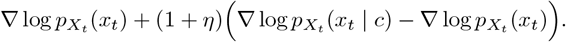

While DDPM and its deterministic variant, the Denoising Diffusion Implicit Model (DDIM) algorithm, achieve remarkable empirical successes, they often suffer from low sampling speed due to extensive function evaluations needed during the sampling phase. In the implementation of our method, we adopt a training-free acceleration scheme that comes with provably faster speed than the original DDPM and DDIM samplers. More details of this acceleration scheme can be found in Li et al. (2024) and other references (see, e.g. Lu et al. (2022); Li et al. (2025)).

### 2.4 Embedding for perturbation

We evaluated generative methods using two genome-scale CRISPR Perturb-seq datasets. The *Yao dataset* assesses the impact of 598 immune-related genes that were knocked out in a human macrophage cell line treated with bacterial lipopolysaccharide (Yao et al., 2024). The study investigated the performance of two experimental procedures, but we restricted our analysis to data derived by conventional perturb-seq methods. The *Replogle dataset* perturbed 2,393 essential target genes in cells of the retinal pigment epithelial cell line (RPE1) using CRISPRi (Replogle et al., 2022). The datasets represent two types of studies that are often illustrated in the literature. The perturbations in the Yao dataset are of genes known to be related to immune function, and hence, we anticipate similar function in many of the perturbations. At the same time, the effect sizes of these perturbations are relatively small. By contrast, the perturbations in the Replogle dataset are of essential genes across the entire genome. The effect sizes of these perturbations are large, and therefore, it should be easier to generate realistic data in this scenario (Figure S1).

Existing methods designed to predict the outcome of unmeasured perturbations utilize various AI tools to produce embeddings for unmeasured perturbations. GEARS uses a graphical neural network based on GO terms and observed gene correlations; scGPT uses a foundation model; and scLAMBDA uses information from an LLM. For our dbDiffusion model, we use simple, data-driven tools to determine the embedding of unmeasured perturbations, as depicted in Figure 3. We have two sources of information: the estimated effect size for each gene by perturbation combination; and the gene expression correlation matrix. From the former, we can cluster perturbations to identify those experiments with similar outcomes (Figure 3a, b). Naturally, the perturbation of interest *K* is not available for this display, but we aim to identify the cluster membership of a perturbation *K* by examining the gene expression correlation matrix (Figure 3c, d).

**Figure 3.**
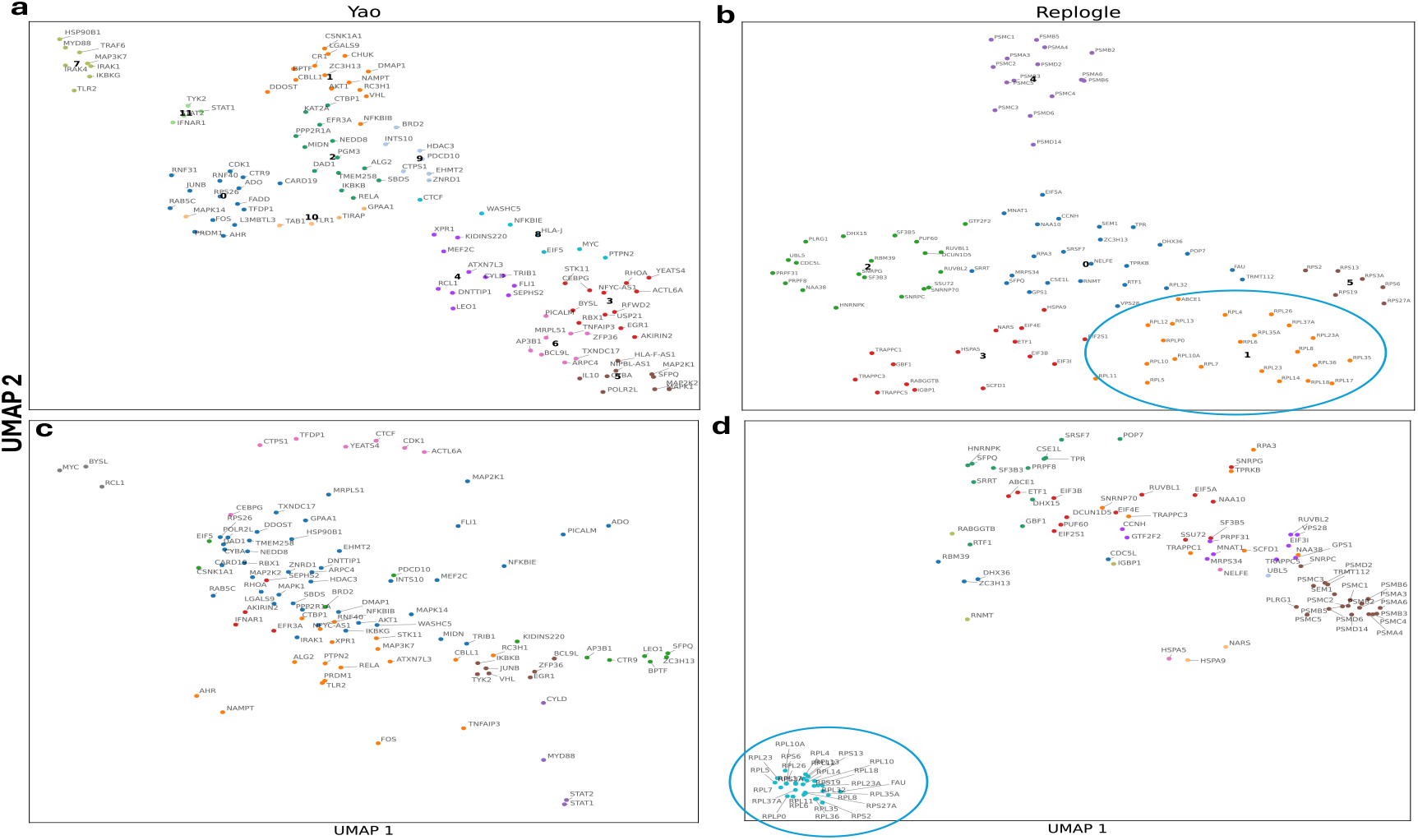
**a. b.** Illustration of clusters of perturbations in (Yao et al., 2024) and (Replogle et al., 2022) respectively. **c. d**. Illustration of clusters of genes in (Yao et al., 2024) and (Replogle et al., 2022) respectively. See Methods for more details.

### 2.5 Debiasing

Debiasing is a key step in our procedure. It is based on the conjecture that the bias in estimating the mean gene expression of a perturbation tends to be similar across the perturbations in a cluster. Consequently, we debias our predicted means, using the observed bias in the other perturbations within the selected cluster. For illustration, we depict the bias associated with the ten genes with the greatest effects of the ZC3H13 perturbation, which we assign to cluster 1 in the Yao dataset. We contrast the realized bias for perturbations in clusters 1 with cluster 5, chosen because it is well separated from ZC3H13. In practice, the true bias for ZC3H13 would not be available, but here we plot this value to reveal how the average bias in cluster 1 estimates the desired bias fairly well, but very poorly in cluster 5 (Figure 4a). We repeat this experiment for the PSMA4 perturbation from the Replogle dataset with similar results (Figure 4b).

**Figure 4.**
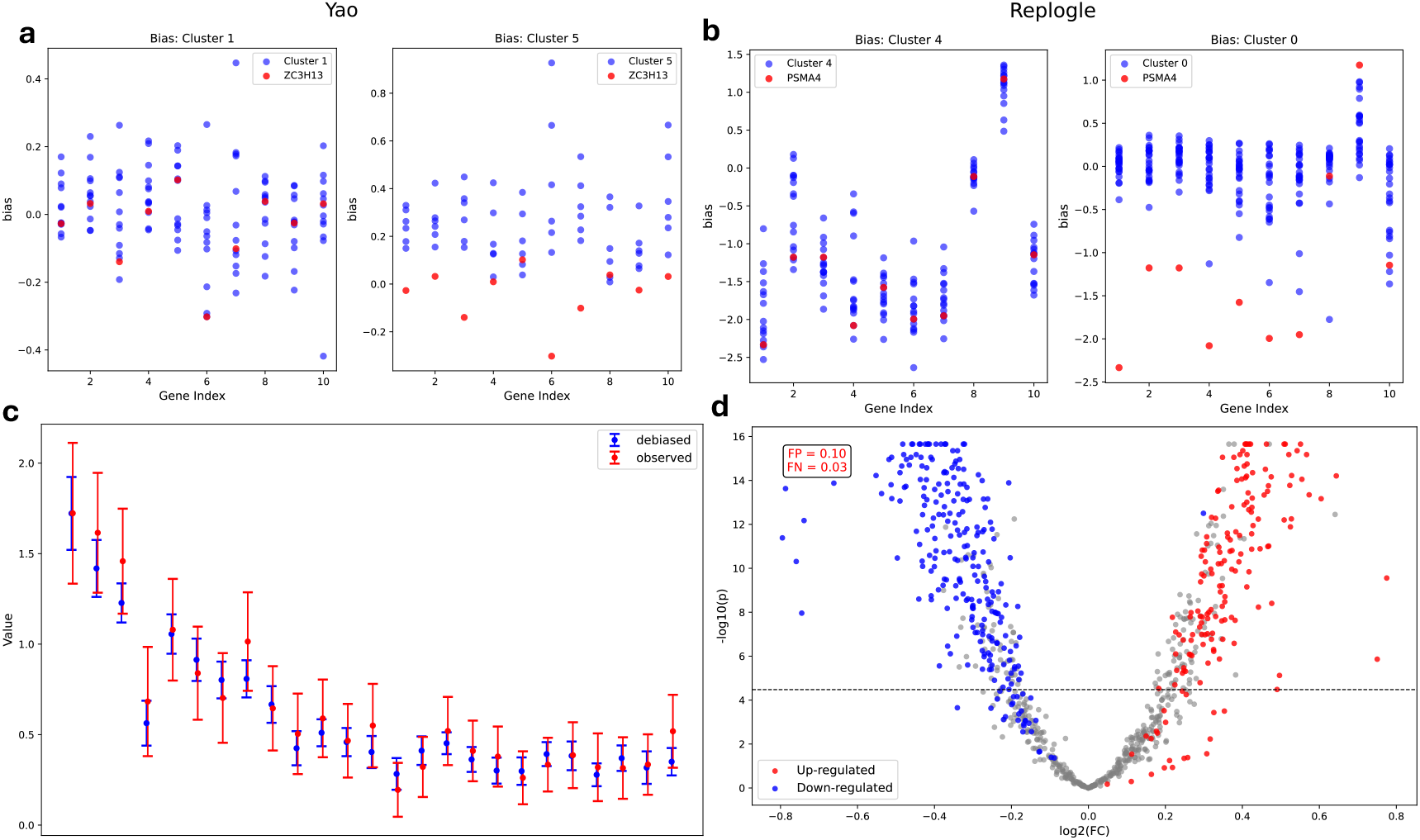
**a**. Biases calculated among all perturbations in Cluster 1 and Cluster 5 on the genes with the biggest effect sizes of ZC3H13. The red dots are calculated when we use the embedding of ZC3H13 without perturbation mapping. **b**. Biases are calculated similarly in the Replogle dataset. **c**. Comparison between the confidence intervals of debiased ones and observed ones. All confidence intervals are constructed for perturbation ZC3H13 of the Yao dataset. **d**. The volcano plot on perturbation PSMA4 in the Replogle dataset. We color dots in grey when genes are calculated using observed z-scores as insignificant. Red dots denote cases where genes are upregulated in the observed data, and blue dots indicate genes that are downregulated. The x-axis shows the difference between the debiased mean and the mean of non-targeting cells, and the y-axis shows the p-values of z-scores of debiased cells.

To illustrate how well dbDiffusion estimates the mean gene expression for an unseen perturbation, we select 25 genes with the largest effect due to perturbation ZC3H13 of the Yao dataset. A comparison between the confidence intervals of the estimated effects and observed ones show substantial overlap (Figure 4c).

Our objective is to predict which genes will be significantly impacted by an unmeasured perturbation. For one perturbation, PSMB5 from the Replogle dataset, we simulate this situation by constructing a test using the generated mean divided by the standard deviation (Figure 4d). Examining the correspondence of the blue (red) points with the significantly down (up) regulated points in the display, it is clear that the generated means and standard errors predict the observed outcomes fairly successfully: the false positive rate is 10% and the false negative rate is 3%.

### 2.6 Benchmarking

We measured the similarity between predicted and observed mean changes in gene expression across test perturbations, including the 14 with the largest effect sizes in Yao’s dataset and the 20 with the largest effect sizes in Replogle’s dataset, using Pearson’s correlation coefficient (PCC, Figure 5a,b). dbDiffusion performed equivalently to cfDiffusion in the Yao dataset and much better in the Replogle dataset, indicating that cfDiffusion’s one-hot encoding model was not able to make efficient use of the data from the targeted perturbation, which was included for training. The results suggest that with a classifier-free diffusion model, an informative embedding can greatly improve performance, even beyond the information gleaned by including the perturbation data for training.

**Figure 5.**
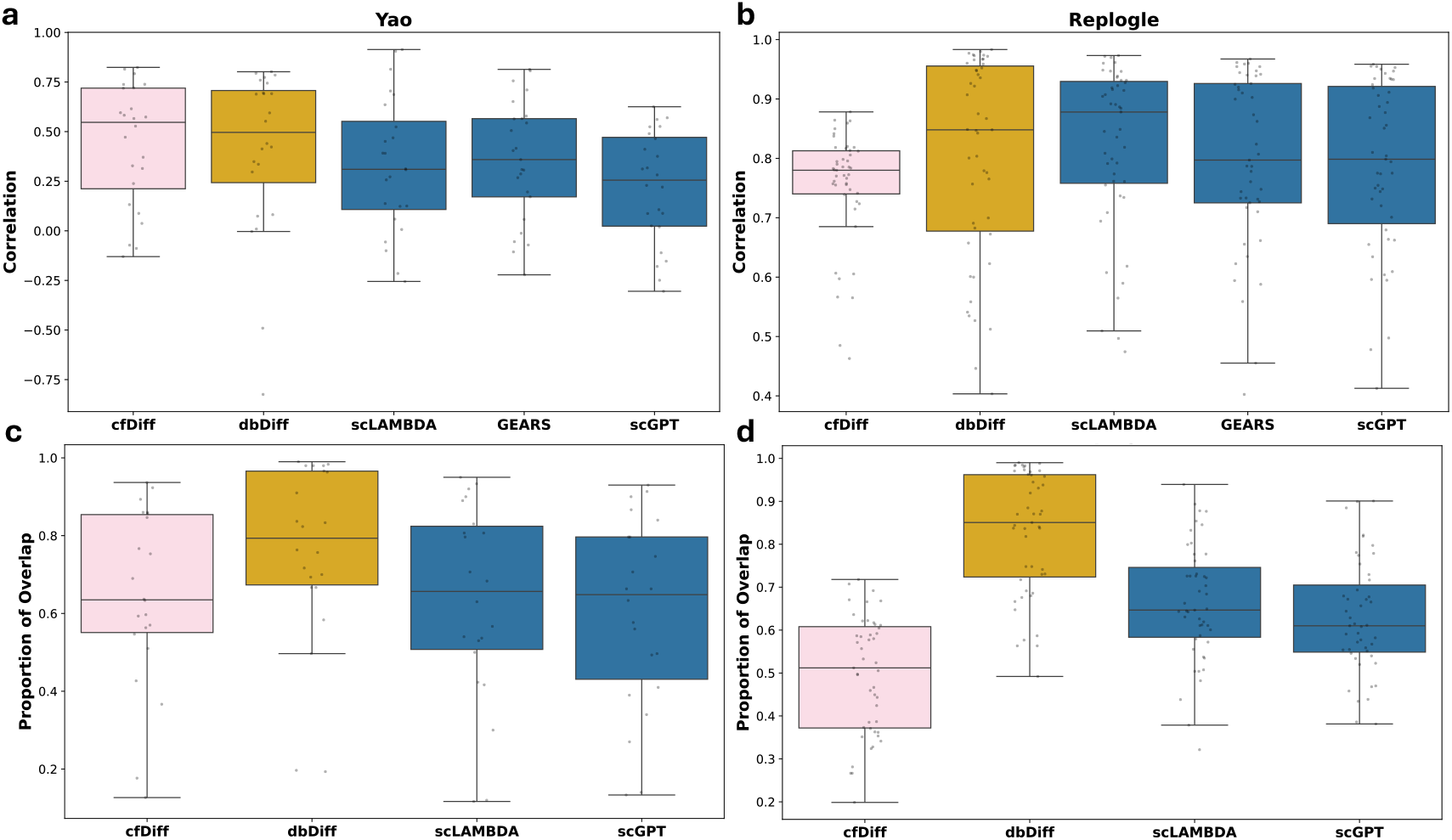
Comparison of Pearson correlation coefficient (PCC) and proportion of overlap in confidence intervals of debiased mean and observed mean for single-gene perturbation outcomes using two datasets: Yao (Yao et al., 2024) and Replogle (Replogle et al., 2022). **a, b.** PCC between the predicted and true average changes in gene expression, evaluated using 300 and 1500 highly variable genes, respectively, across different methods. **c, d**. Proportion of overlap in confidence intervals of debiased mean and observed mean, evaluated using 300 and 1500 highly variable genes, respectively, across different methods.

Regarding benchmark comparisons, dbDiffusion outperformed the other three methods in the Yao dataset, achieving an average PCC of ∼ 0.5, considerably higher than the next competitor (Figure 5a, b). For the Replogle dataset, the performance was slightly inferior to that of scLAMBDA but much better than that of GEARS and scGPT. The performance of all methods was considerably lower in the Yao dataset than in the Replogle dataset. We expect that this is due to the fact that the effect size of the assessed perturbations is smaller in the Yao dataset (Figure S1).

We are also interested in how well a confidence interval for the mean effect overlaps with the truth. A high overlap supports our ability to make inferences about the impact of an unseen perturbation. dbDiffusion clearly has the highest overlap, followed by scLAMBDA (Figure 5c, d). It is not possible to assess GEARS by this criterion because it does not generate gene by cell counts and hence does not support estimation of a confidence interval.

The performance of dbDiffusion benefits from the debiasing step. To investigate whether this concept could be helpful for other methods, we computed the bias of each estimator using the cluster of perturbations estimated by our model. Then we subtracted the bias directly from the predicted mean expression for models cfDiffusion, scLAMBDA, and scGPT, and each improved considerably (Figure 6). Naturally, these improvements required input from the clustering model that is the basis of dbDiffusion, so they are not valid comparisons of methods. However, the improvement suggests that this type of modification applied to other methods could lead to better performance.

**Figure 6.**
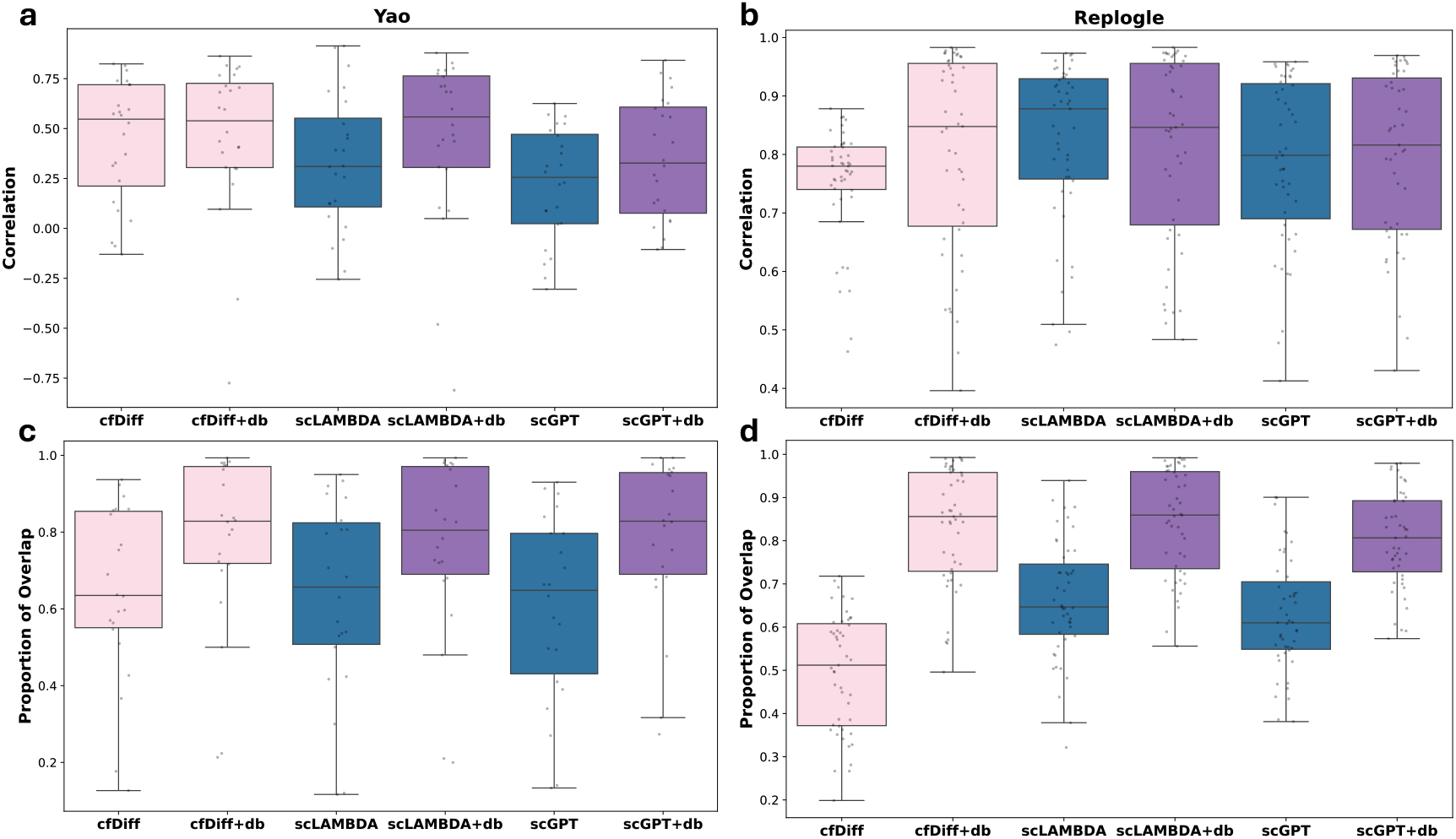
**Comparison of Pearson correlation coefficient (PCC) and proportion of overlap in confidence intervals of debiased mean and observed mean for single-gene perturbation out-comes using two datasets: Yao (Yao et al., 2024) and Replogle (Replogle et al., 2022), across different methods along with the debiased versions. Perturbations and genes evaluated are the same as Figure 5**

### 2.7 Biological insights

The performance of the estimators varies considerably between different perturbations, with some perturbations producing much lower PCC than others for a given method (Figure 5). We aim to determine whether some perturbations are inherently hard to predict or if our method is challenged by particular perturbations that are less difficult for other methods and vice versa. Starting with the profiles of predicted mean change in gene expression for each perturbation, relative to control, we examine the pairwise correlation between methods for each perturbation. We find that the resulting estimators are highly correlated across all perturbations for the Replogle dataset, but weakly correlated in the Yao dataset (Figure 7a). This supports our conjecture that the signal in the Replogle dataset is strong enough so that every method performs well. By contrast, the signal in the Yao dataset is much weaker, and therefore the methods struggle to predict the outcome of these perturbations (Figure S1).

**Figure 7.**
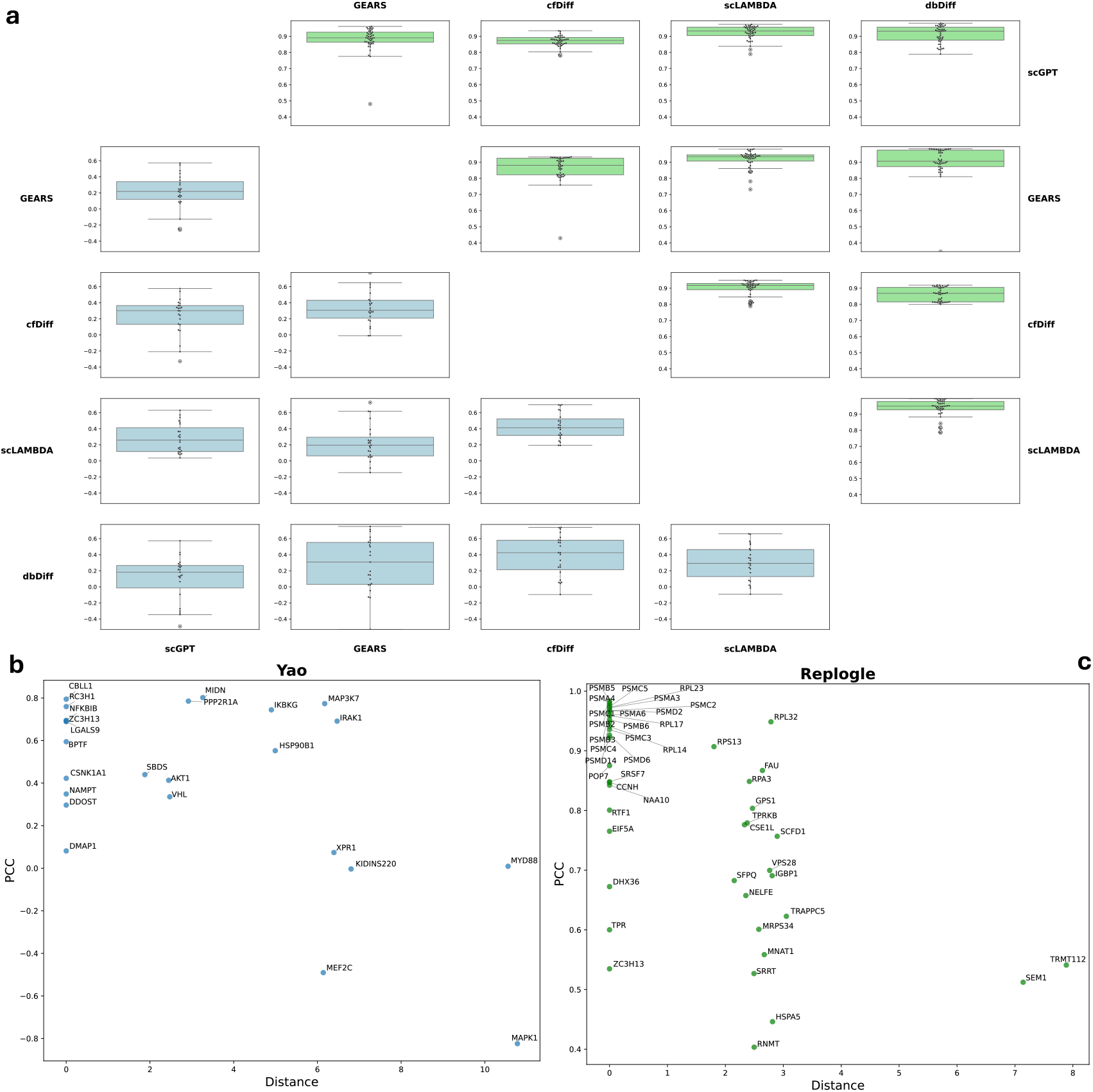
a. Pair-wise comparison of different methods of gene-level prediction correlation. For each pair of methods, we calculated the mean of gene-level predictions minus that of the non-targeting cells, respectively. The correlation between these two sets of means is then computed for each perturbation, yielding one correlation value per perturbation. Each boxplot in the figure corresponds to one pair of methods, and each dot within the boxplot represents the correlation value for a single perturbation. **b. c**. The relationship between PCC and the distance between the cluster assigned to the perturbation evaluated and the cluster of the ground truth.

Next, we aim to determine what properties render some perturbations more difficult than others. We begin by investigating the performance of dbDiffusion across perturbations. Many perturbations can be mapped back to their appropriate cluster using our algorithm, but sometimes this algorithm fails, and the embedding places the perturbation in a cluster a substantial distance from the optimal one. In that case, the quality of the predictions declines substantially (Figure 7b, c). To understand this phenomenon, we examine several variables to determine factors that influence PCC and distance from the optimal cluster.

A likely factor determining success is the similarity of genes between the effect size and gene expression clusters. A perturbation can only be mapped to the appropriate effect size cluster if those same genes are clustered due to correlated gene expression. To measure this feature, for each perturbation, we identify its cluster based on effect size, then examine the distribution of these genes across the gene expression clusters. To quantify the results, we borrow an idea from Rand Index and count the number of pairs of genes that appear in the same gene-expression cluster, relative to the total number of pairs, and call the measure RI (see Supplemental Information).

Finally, we explore two models. One to predict the distance of the estimated cluster from the optimal cluster, and another to predict the achieved correlation of the generated perturbation (PCC) (Figure S1. a). Predictors include the average effect size, sample size, RI, and measures of connectedness of both the effect size and the gene expression clusters. When predicting PCC, the distance is strongly negatively correlated (*r* = -.69). And when predicting distance, RI is strongly negatively correlated (*r* = -.44). No other variables are predictive for either response variable. We conclude that dbDiffusion is most successful when the clusters in effect size and gene expression space are similar. This characteristic largely determines the distance. A poorly matched clustering leads to a poor embedding and ultimately a poor estimate of the response to perturbation.

To see these relationships in action, consider three of the Yao dataset’s perturbations in more detail (Figure S1. b, c). dbDiffusion performs very poorly for MEF2C, but each of the other methods performs acceptably. From Figure 7b it is clear that this perturbation was very poorly placed in the embedding scheme and consequently the predictions were poor, likely due to poor clustering (RI = 0.81, a fairly low value).

Next, consider AKT1. For this perturbation, we obtain a suboptimal but acceptable embedding partly due to good clustering (RI = 1.0, a fairly high value). All other methods perform poorly for this perturbation. To determine their embeddings, each of these methods relies on external information via AI methods: scGPT relies primarily on a large foundation model; scLAMBDA relies on LLMs trained using publications in NCBI; and GEARS derives embeddings from networks of GO terms, as well as the correlation of gene expression. This is an example where the AI input is not well aligned with the experiment.

Finally, consider RC3H1. For this perturbation, we obtain the optimal embedding likely due to good clustering (RI = 1.0). Although the performance of other methods is lower than that of dbDiffusion, scLAMBDA and GEARS provide acceptable predictions. scGPT does not. Presumably, this is because the foundation model was biased relative to the conditions of this particular experiment.

In practice, it is challenging to know which method will perform best for a particular perturbation; however, we do have some options for calibrating the performance of dbDiffusion. Although it is not possible to determine whether a particular unmeasured perturbation will have a high RI, it is possible to assess the RIs of related perturbations. The RIs of genes that are highly correlated with the perturbation of interest in the gene expression space should be predictive. Moreover, one can use AI tools, such as LLMs and GO terms, to identify measured perturbations that are expected to have a similar biological function. Then evaluate the RIs of these perturbations. By contrast, it is much more difficult to predict when methods based exclusively on LLMs and Foundation models will perform poorly.

## 3 Discussion

In this work, we introduced dbDiffusion, a diffusion-based generative framework for predicting single-cell responses to unseen perturbations. By integrating perturbation embeddings with classifier-free guidance and, critically, incorporating a debiasing step, our method consistently outperformed state-of-the-art approaches such as GEARS, scLAMBDA, and scGPT across benchmark Perturb-seq datasets. dbDiffusion demonstrated particular strength in accurately estimating mean gene expression effects, constructing valid confidence intervals, and mitigating systematic bias in generative predictions. Importantly, performance varied by dataset: the method excelled in the Replogle dataset, where perturbation effects were large, while predictions in the Yao dataset proved more challenging due to smaller effect sizes. These findings highlight both the strengths and limitations of generative perturbation prediction.

Our study further revealed that predictive accuracy strongly depends on how embeddings are constructed. Data-driven embeddings based on clustering of effect sizes and correlated gene expression were especially effective, particularly when clusters in both spaces aligned closely (high RI values). In contrast, embeddings derived solely from AI-based tools (e.g., LLM-informed scLAMBDA or foundation model embeddings in scGPT) occasionally provided complementary advantages when biological clustering was weak, but they also introduced biases. These results suggest that hybrid or adaptive embedding strategies could provide the best of both approaches.

A central challenge lies in anticipating, for a new perturbation, which predictive framework will be most reliable. For dbDiffusion, embedding quality emerged as the dominant factor: when a perturbation mapped to a well-matched cluster, performance was strong; when cluster alignment was poor, alternative methods sometimes fared better. The Rand Index (RI), which quantifies the consistency between effect-size and gene-expression clusters, proved a valuable predictor of success. High RI values corresponded to accurate predictions, whereas low RI values signaled a higher risk of error. Thus, a practical strategy is to evaluate the RIs of related measured perturbations before applying a method to an unseen perturbation. If RI is high, dbDiffusion should be favored; if low, complementary embeddings (e.g., from LLMs or GO networks) may provide more reliable guidance.

Compared to purely AI-based tools, which often lack interpretability and exhibit limited generalization, our approach is grounded in statistical intuition and remains amenable to model diagnosis. More broadly, these findings point toward the utility of ensemble or meta-predictor frameworks, in which the choice of predictive model is dynamically guided by dataset properties such as effect size, cluster quality, and embedding consistency.

## 4 Methods

### 4.1 Data preparation

For the Replogle dataset, we calculated the effect size matrix using the SCEPTRE software Barry et al. (2024). For the Yao dataset, we downloaded the effect size matrix computed using similar methods Yao et al. (2024). The Yao’s dataset reports response to 599 perturbations for 16,952 genes, while the Replogle’s dataset reports response to 2393 perturbations for 8,748 genes. Prior to analysis, we reduced the gene and perturbation lists to focus on more informative outcomes. We first select ∽100 perturbations with the highest total absolute effect sizes across all genes. We then restrict the effect size matrix to these selected perturbations. Next, we identify around 4,000 genes with the largest total absolute effect sizes across the chosen perturbations. These genes are all highly expressed, so to further select informative genes, we subset the gene expression matrix to the same about ∼100 perturbations and then choose the 1,500 genes with the highest total expression values. Ultimately, for both datasets, the effect size matrix and the gene expression matrix used in our analysis include 1,500 genes.

The software takes as input the UMI counts, which are transformed internally. All other analysis is based on the standard log transformation, as used in Seurat: UMI counts are divided by total counts for the cell, multiplied by 10,000, and then subjected to a log(*x* + 1) transformation.

### 4.2 Identifying the Most Similar Perturbations for Unseen Perturbations Using Two-Step Clustering

Before running the generative model to sample the unobserved (or target) perturbation, we first need to determine its embedding, which serves as input to the diffusion model during the sampling process. Similarly to how we obtained the embedding for an unobserved tissue in Figure 2d, we first identify the perturbations most similar to the unobserved. The embedding of the unobserved perturbation is then set as the mean of the embeddings of these identified perturbations. To achieve this, we apply a two-step Leiden clustering Traag et al. (2019) as follows:

We first perform PCA on the effect size matrix, followed by Leiden clustering to group the perturbations into effect-size clusters, denoted as 𝒞_*ES*_ ={ *P*_1_, …, *P*_*E*_}, where *P*_*i*_ represents a cluster of perturbations.

Next, given the ℝ^*n×p*^ gene expression matrix for all cells (excluding the perturbation of interest), we approximate the gene correlation matrix, retaining the *K*_1_ top eigenvectors. From this approximate correlation matrix, we obtain more smoothness, using the top *K*_2_ eigenvectors. We then apply Leiden clustering on this smoothed representation of the correlation to obtain gene-expression clusters, denoted as 𝒞_*GE*_ = {*G*_1_, …, *G*_*F*_}, where *G*_*i*_ represents a gene cluster.

Finally, for simplicity of exposition, we assume that the unobserved perturbation affects the gene that falls in *G*_1_. For each *i* = 1, …, *E*, we calculate the number of perturbations in *P*_*i*_ that overlap with *G*_1_: *n*_*i*_ := |*P*_*i*_ ∩ *G*_1_|, and the proportion of matched perturbations: 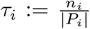. Sometimes, choosing the “best” cluster is challenging—for example, if *n*_*i*_ *> n*_*j*_ but *τ*_*i*_ *< τ*_*j*_, then *P*_*i*_ contains more matched perturbations but is also much larger, potentially introducing noise into the embedding estimate (illustrated in Figure 8).

**Figure 8.**
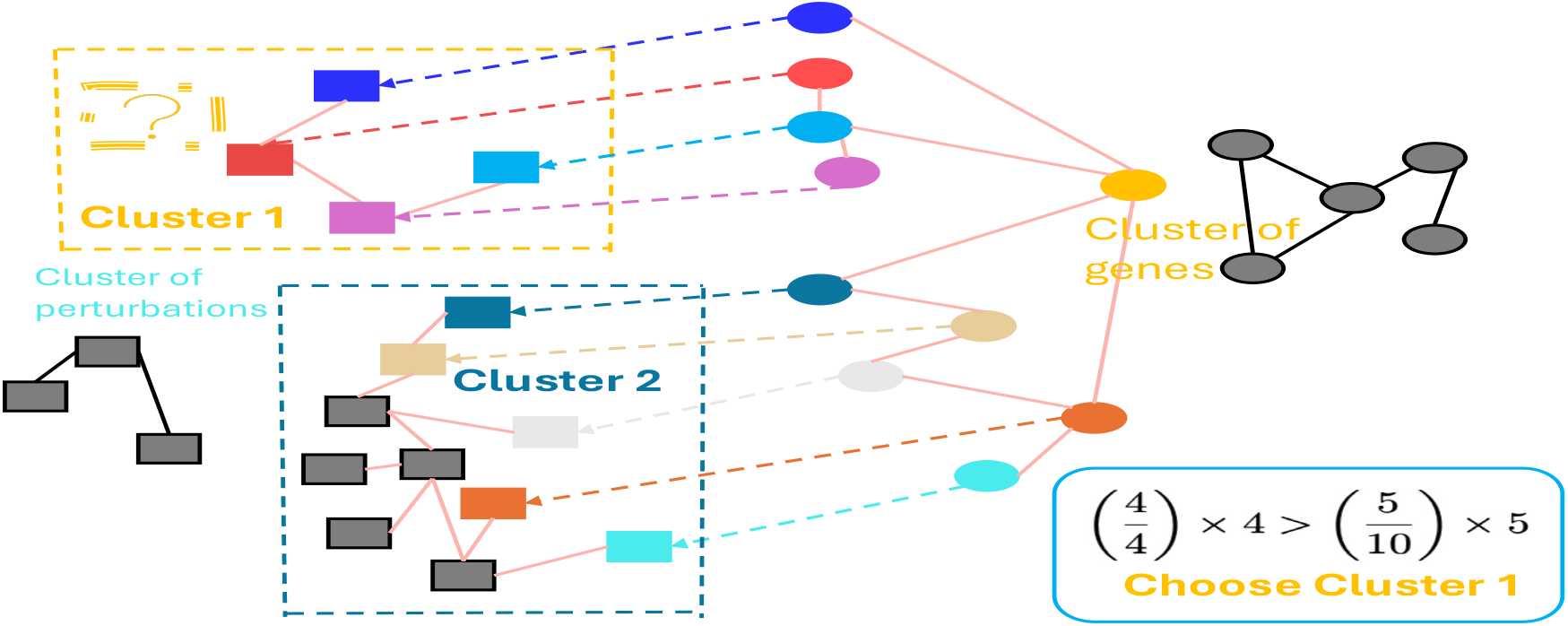
Illustration of assigning the perturbation unobserved to a cluster of perturbations observed.

To balance the number of matches and the proportion, one can use a user-defined parameter *β >* 0 and define a score for each cluster: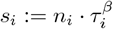. The selected cluster is then

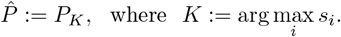

The embedding of the unobserved perturbation is set as the mean embedding of the perturbations in 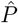.

### 4.3 Debiasing the Estimator

After running the generative model, we will analyze each outcome gene individually so that *Y*_*ijk*_ is the expression of the *i*’th observation, *j*’th gene from perturbation *k*. We index the perturbations with a vector *X*_*k*_ that indicates the embedding of the perturbation that the cell has received *k* = 1, …, *K* − 1. *K* indicates the perturbation of interest that has not been measured.

Our goal is to estimate *E*[*Y*_*j*_|*X*_*k*_] and test whether this differs from the mean of the control sample. If we had *n* actual measurements for *K* (*Y*_*ijK*_), matched predictions (*Ŷ*_*ijK*_), and *N* new predicted measurements 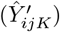, then the PPI procedure (Angelopoulos et al., 2023) would be as follows:

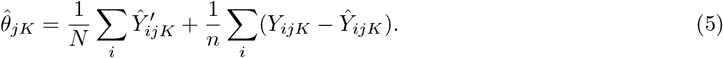

The estimator shares features with the well-known semiparametric debiased estimator, where the second term plays the role of a bias term. There are three big differences between the scenario above and ours. First, we do not have a prediction model to generate gene expression, given a set of cellular-level covariates. Rather, we generate an entire set of observations under a particular diffusion model. Hence, there is no natural analog to *Ŷ*_*ijK*_, which pairs with *Y*_*ijk*_. Second, in our situation, we have no measurements for the *K*’th perturbation, which would play the role of *Y*_*ijK*_. What we do have is a gene embedding+diffusion model to predict *N* cells that have the new perturbation 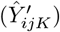.

Our biggest challenge is how to estimate the bias when we do not have an available analog for the second term above. We can not assume that the bias is common across all perturbations. Instead, we cluster observed perturbations, based on our gene embedding model, and estimate the bias based on these perturbations. We assume that the bias is well estimated by averaging the observed bias for *K*_*c*_ perturbations in cluster 𝒞_*K*_. With this assumption, we can modify the PPI estimator for our purpose.

For each observed perturbation in 𝒞_*K*_, set aside these data, sequentially, fit the diffusion model, and predict the values 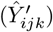 for *N* cells. Likewise, predict *N* observations of the unobserved perturbation 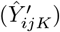. Using these quantities, we define our estimator as follows:

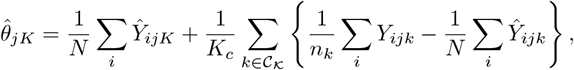

where *Y*_*ijk*_ is the original data under perturbation *k* with sample size *n*_*k*_.

To analyze our estimator, take the expectation of each term above. We find that 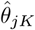 is consistent under two scenarios: (1) naively, if the true impact of the *K*’th perturbation is equal to the average effect in cluster 𝒞_*K*_ then the estimator is trivially consistent; or (2) if bias in the generated data for the *K*’th perturbation equals average bias in cluster 𝒞_*K*_, the estimator is consistent.

Assuming that *N* is much larger than *n*, it follows that 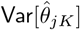 is well approximated by the variance of the second term. Formulating a model for the effect of perturbation *k* on gene *j*, we write

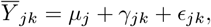

where *µ*_*j*_ is the effect of the gene, *γ*_*jk*_ is the effect of the perturbation with variance 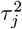 within the cluster, and *ϵ*_*jk*_ is the variability of the mean measurement error with variance 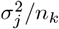. It follows that

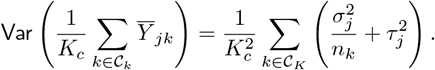

A natural estimator for 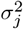 is

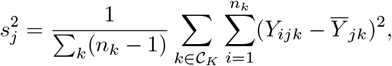

and for 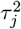 is

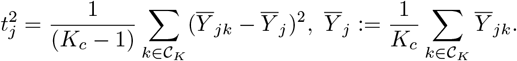

In total, the standard error can be estimated as

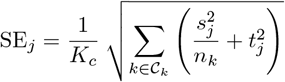

We obtain an approximately (1 − *α*)% confidence interval as

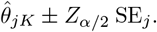

To test each gene for an effect due to the perturbation *K*, we use

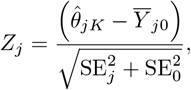

where the zero subscript indicates the control sample, with sample size *n*_0_, and 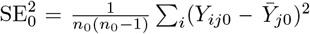.

## 5 Ackowledgement

We thank two referees for helpful suggestions. This work was funded, in part, by a grant to the NIMH (MH123184). Y. Wei is supported in part by the NSF grants CCF-2418156 and CAREER award DMS-2143215.

## A. Supplemental Information

### A.1 Correlation Analysis

We formulate a model for the mean for gene *j*, perturbation *k* (*k* = 0, for control) and method *ℓ* (*ℓ* = 0, for observed data). The model for the mean response includes a random effect for gene, perturbation, and bias *b* of the method:

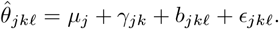

The observed perturbation is

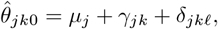

and the control mean is

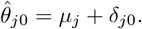

Terms in the model are random effects, with variance notation 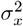 where *x* represents the source of variability. All these terms are assumed to be independent, except for the covariance between the perturbation effect and the bias in the estimator: *Cov*(*γ, b*) = *σ*_*γ,b*_. The most likely bias of a method is towards the control, that is, *b*_*jkℓ*_ *∝* −*γ*_*jk*_. Consequently, we expect the covariance term to be negative.

The correlation between the estimated mean, 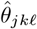, and the observed mean, 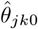, over genes will be dominated by 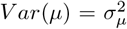, assuming that the genes vary substantially in their expression. Consequently, the correlation will be near 1. To circumvent this problem, center each mean by the control mean, i.e., let

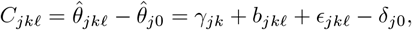

and, for the observed perturbation, let

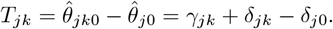

It follows that

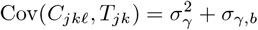

Notice that this means that the correlation of these centered quantities will be zero unless there is some variability due to perturbations.

Now, assuming there is a perturbation effect, we need to look at the correlation to see differences in performance.

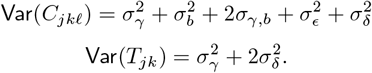

The correlation of a method can be reduced by having a larger error (*σ*^2^), but is more likely to be diminished due to a relatively large bias. And

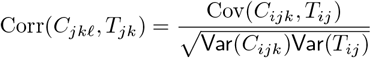

The Corr(*C*_*ijk*_, *T*_*ij*_) can be near zero or even negative if the variability due to bias and estimation errors is large. Our dbDiffusion estimator aims to remove the bias, whereas no other method has this feature. The good performance of the debias function relies on our ability to choose observed perturbations that are similar to the unobserved one. We can then use the bias of estimation at these perturbations to correct the diffusion estimator.

To compare estimators from a pair of methods, we can examine the vectors *C*_*jkℓ*_, *j* = 1, …, *G* for a fixed *k*. For each pair of methods, compute the pairwise correlation of the centered estimates. This correlation will be big if two methods produce similar estimates at a per-gene/perturbation level.

**Fig. S1.**
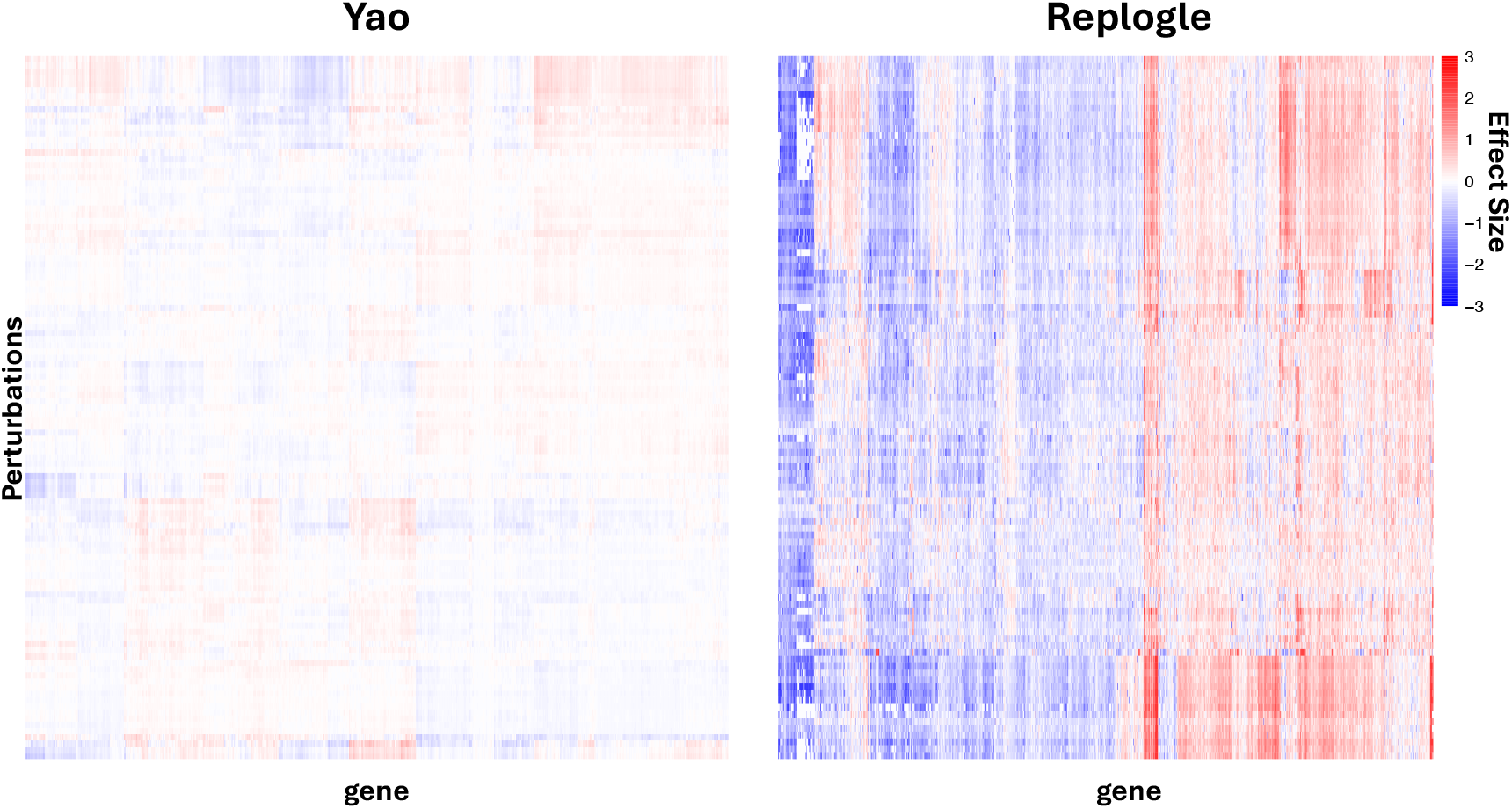
Heatmaps of Effect sizes in the Yao and Replogle datasets.

### A.2 Definition of Measure Random Index

We consider the target perturbation, denoted as *p*_1_, and the group calculated by the embedding matrix(effect size matrix) of size *n* that contains *p*_1_: *S* ={ *p*_1_, …, *p*_*n*_}. It is direct that in the cluster by embedding matrix, all *n* perturbations are gathered in the same cluster. However, in the cluster calculated by the gene expression matrix, which surely do not contain cells under *p*_1_, the clusters in which different perturbations *p*_*i*_, *i* = 1, …, *n* fall are different, denoted as *n* labels for each perturbation *p*_*i*_: **L** = {*L*_1_, …, *L*_*n*_}. As a result, we can count the number of **distinctions** in 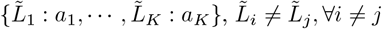, where *a*_*i*_ shows the appearing times of 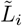 in **L** such that 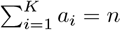. The random index (RI) is calculated as

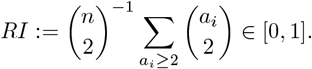

Intuitively, when all *L*_*i*_ are the same, two types of clusters share the highest consistency and *RI* reaches the maximum 1. As another extreme, when *L*_*i*_ ≠ *L*_*j*_, *∀i* ≠ *j*, the cluster by the gene expression matrix shows no consistency with the one by the embedding matrix, since all perturbations we evaluate fall into the same cluster by the embedding matrix, then *RI* reaches the minimum 0.

**Fig. S2.**
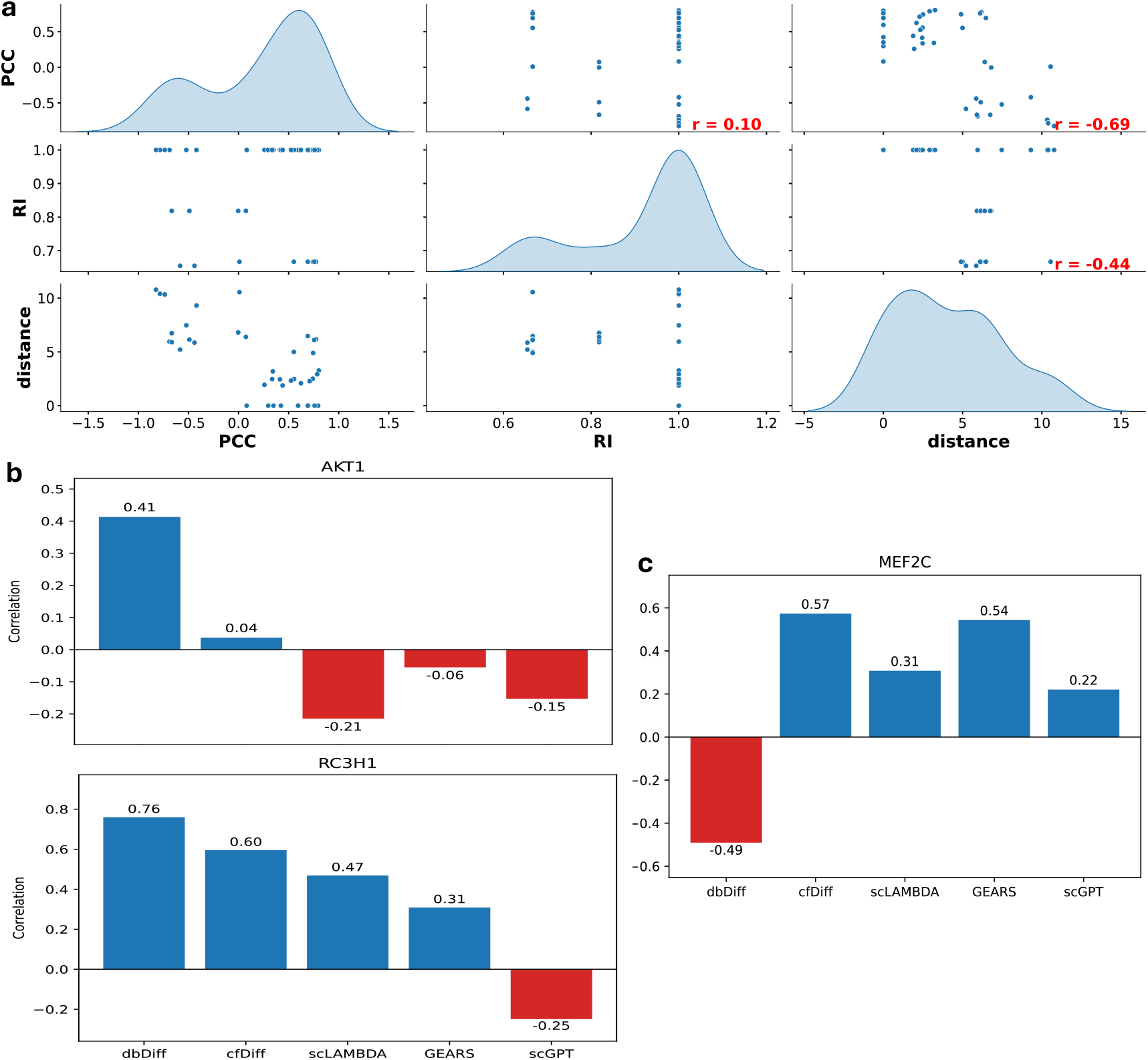
**a.** Scatterplot showing the relationship between PCC and RI, PCC and distance between the cluster assignment by our algorithm and the ground truth cluster, distance, and RI, with pairwise correlation indicated. **b.** PCC of perturbations AKT1 and RC3H1 across different prediction methods, where dbDiff performs well, while others do not perform so well. **c.** PCC of perturbation MEF2C across different methods, where dbDiff performs badly while others perform well.

## Footnotes

1 GitHub at: https://github.com/ergan-shang/dbDiffusion.

